# Discrete confidence levels revealed by sequential decisions

**DOI:** 10.1101/169243

**Authors:** Matteo Lisi, Gianluigi Mongillo, Andrei Gorea

## Abstract

Whether humans are optimal decision makers is still a debated issue in the realm of perceptual decisions. Taking advantage of the direct link between an optimal decision-making and the confidence in that decision, we offer a new dual-decisions method of inferring such confidence without asking for its explicit valuation. Our method circumvents the well-known miscalibration issue with explicit confidence reports as well as the specification of the cost-function required by ‘opt-out’ or post-decision wagering methods. We show that observers’ inferred confidence in their first decision and its use in a subsequent decision (conditioned upon the correctness of the first) fall short of both the ideal Bayesian strategy, as well as of an under-sampling approximation or a modified Bayesian strategy augmented with an additional bias term to accommodate global miscalibration of confidence. The observed data are instead significantly better fitted by a model positing that observers use only few confidence levels or states, at odds with the continuous confidence function of stimulus level prescribed by a normative behavior. These findings question the validity of normative-Bayesian accounts of subjective confidence and metaperceptual judgments.

## Introduction

How do we select behavioral responses when interacting with a given environment? According to a popular theory - the rational choice theory - human agents evaluate, for each decision to be taken, the expected utility of each possible course of action, and select thereafter the one yielding the highest rank. Despite the fact that the *descriptive* adequacy of rational choice theory has long been challenged on empirical as well as on theoretical grounds, mainly questioning its biological/psychological plausibility [16,17,20,29,42], the debate is far from settled [5,6,19]. In order to maximize expected utility, an agent has to be able to associate a *subjective* probability to each possible consequence of its actions. The characterization of this process requires (i) the quantitative assessment of the agent’s ability to attach probabilities to events, and (ii) the appraisal of the measured subjective probabilities against those of a Bayesian observer, with the same prior knowledge as the agent, would assign to the same events. The experimental instantiation of such comparisons has been hindered by serious methodological problems with assessing subjective probabilities.

Subjective probability is tantamount to the agent’s confidence in the occurrence of an event *E* [10]. It is formally defined as a marginal rate of substitution [10,40]: the agent’s subjective probability about the event *E* occurring or having occurred would be *p*(*E*) if the agent is indifferent to gaining one unit of utility contingent on *E* against gaining *p*(*E*) units of utility for sure. Current methods for measuring subjective probabilities in, for example, perceptual decisions (that is, the confidence about choices being correct) using opt-out or post-decision wagering techniques are straightforward operationalizations of the above definition. A major, well and long known problem with these methods is that they rely on unverifiable assumptions about the utility function of the participant (see, e.g., [40]). Such methods cannot disentangle subjective probability from factors such as opportunity cost in waiting-time paradigms. More frequently used, methods requiring *explicit* confidence valuation by the decision-maker suffer from well-documented miscalibrations and response biases (see, e.g., [14,15,28]).

**Figure 1:**
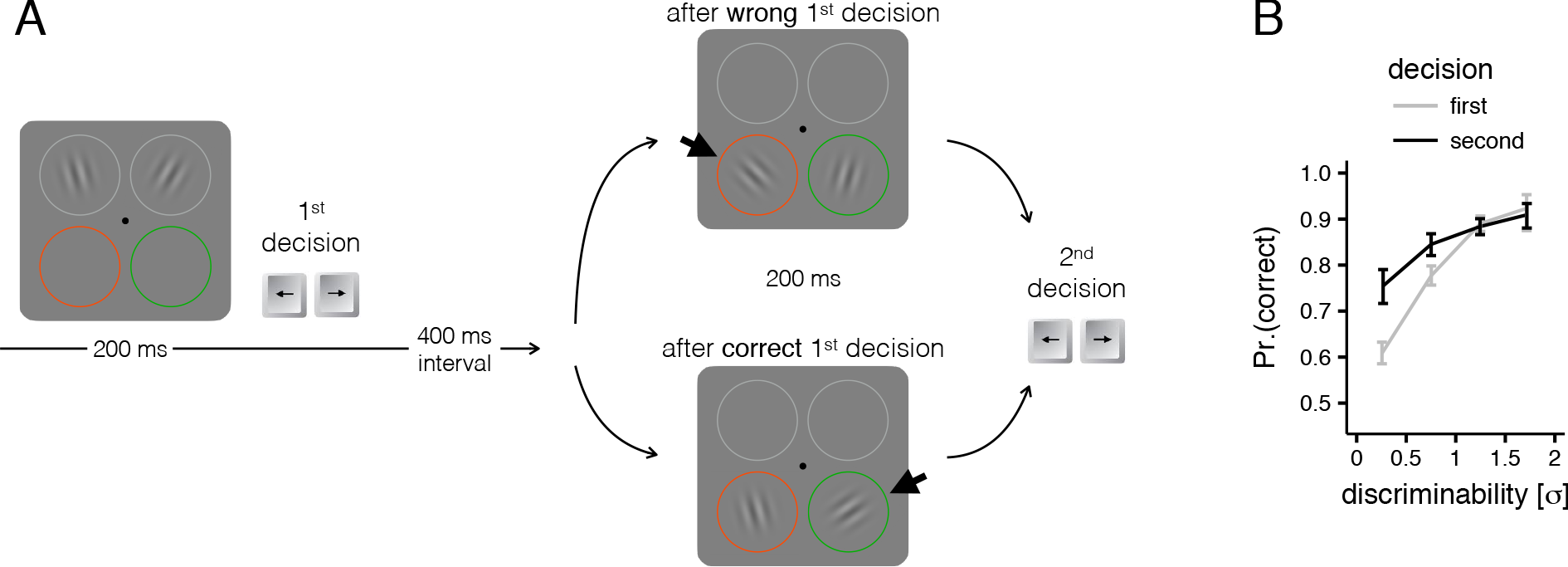
Experimental paradigm and results. **A**. Orientation-orientation (O-O) paradigm. Participants had to make two consecutive decisions on which of two Gabor-patches was more tilted away from the vertical (hence the signal on which the participant had to base the decision was the signed difference in orientation between the two Gabors). While the absolute difference in tilt between the two Gabors (the discriminability) is drawn independently from a uniform distribution in each of the two Gabor-pairs, the location of the more tilted Gabor in the second Gabor-pair (left/right; indicated by the short thick black arrows in the right-hand panel) was made dependent on the accuracy of participant’s first response. **B**. Performance (proportion of correct responses) plotted as a function of discriminability (measured in units of internal noise) separately for 1^st^ and 2^nd^ decisions (averaged over all conditions and experiments). Note how performance in the 2^nd^ decision is higher, especially for trials where the sensory evidence (or discriminability, expressed in standard deviations of internal noise) is small and, in the absence of prior information, performance would be at or near chance. Error bars represents bootstrap 95% CI.

Here, we present a novel approach of estimating subjective probabilities which overcomes the problems above. Human participants were presented with two consecutive signals and asked to decide whether they were above or below some reference value. The key innovation was that the statistics of the second signal was made contingent upon the decision-maker having made a correct decision on the first signal (explicit feedback is not provided on a trial-by-trial basis): correct/incorrect first decisions resulted into signals above/below the reference value for the second decision (a diagram of the experimental protocol is shown in Fig. 1 in the context of an orientation discrimination task; the signal here is the difference in orientation between the two Gabors). Differences in performance between the second and the first decisions, at signal parity, allow the estimation of the subjective probability of being correct on the first decision (i.e., the confidence). Using this approach we show that humans are quite accurate in assessing confidence, yet they exhibit systematic deviations from optimality, such as global under-confidence and high-confidence errors. These systematic deviations cannot be accounted for either by a biased Bayesian observer (which perform Bayesian computations using a biased estimate of the variability of the internal signals) or by a sample-based approximation of the optimal Bayesian strategy (see Supplemental information). An alternative non-Bayesian model, characterized by a finite number of discrete confidence levels, provides the best and most parsimonious description of the empirical patterns of humans’ choices. These results suggest that the evaluation of subjective confidence in human observers is not based on a full posterior distribution, and provide new cues about which kind of shortcuts and approximation may be used by the brain to handle meta-perceptual uncertainty.

## Results

### The dual-decision paradigm

The paradigm is illustrated in Figure 1A with an example of the orientation discrimination task (orientation-orientation or O-O condition, see Material and methods and Supplemental information for details). Participants are presented with two consecutive Gabor-pairs, and for each pair must decide which of the two Gabors was more tilted with respect to the vertical; correct/incorrect first decisions result into displaying the second pair with the more tilted Gabor in the right/left place-holder, respectively, and the participants are told so. The very same experimental format was used with a duration discrimination task (duration-duration or D-D condition) where participants had to decide which of two Gaussian blob flashes (presented sequentially) was displayed for a longer duration. These two conditions were tested both in a first experiment where the difficulties of first and second decision were independently drawn from a uniform distribution (random-pairs experiment) and, on a different group of subjects, in another experiment where these difficulties were not independent (correlated-pairs experiment): more specifically difficult/easy first decisions were more likely to be followed by easy/difficult second decisions, respectively (see Material and methods). This correlation was introduced to encourage participants to rely more on information provided by the first pair of stimuli and exploit the statistical dependence between the signals in the two decision. Additionally, we also tested a condition where the two tasks were combined, duration-orientation or D-O condition, to test within our paradigm the proposal that confidence may work as a ‘common currency’ between different perceptual judgments [12,13].

**Figure 2:**
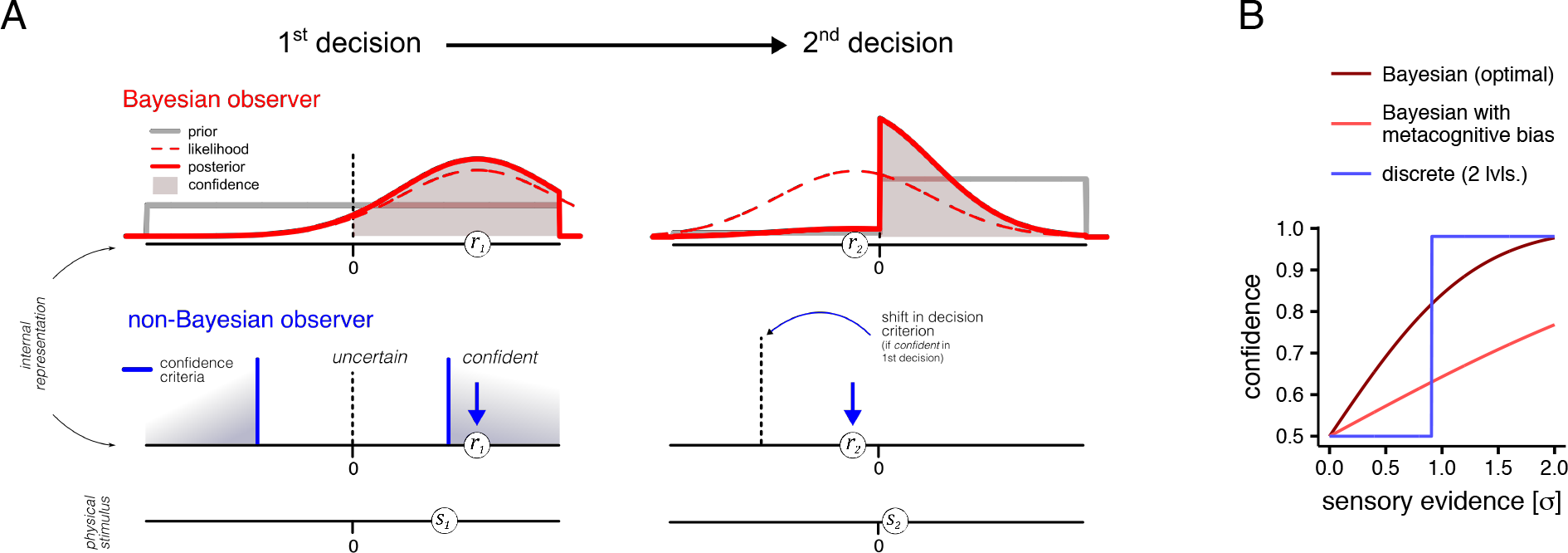
Bayesian and a class of non-Bayesian models of confidence and of sequential decision-making. **A**. Internal beliefs of the Bayesian (top panels) and non-Bayesian observer (bottom panels), for 1^st^ and 2^nd^ decision (from left to right). The lowest abscissae represent the state of the world (e.g. the physical difference in orientation between the two Gabors in the orientation task of Fig. 1) while the internal representations of the decision variable (the sensory evidence) are referred to the upper two abscissae. The first stimulus in this ad-hoc trial (si) evokes a noise contaminated internal response (*r*_1_ = *s*_1_ + *η*, where 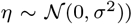. The Bayesian observer (top panels) knows about the statistics of its perceptual system and is able to compute the likelihood function *p*(*r*_1_|*s*_1_). Because for the 1^st^ decision the prior is flat (grey lines in the upper panels), the likelihood (dashed curves) largely overlaps with the posterior (red continuous curves). The Bayesian probability that represents the subjective confidence c_1_ that the true value of the signal lies in the positive semi-axis corresponds to the area of the posterior distribution shaded in red (*c*_1_ = *p*(*ŝ*_1_ > 0|*s*_1_)). The observer knows that the second signal will be drawn from the positive interval (right-hand side) after a correct 1^st^ decision, and from the negative semi-interval after an error. Hence the ideal observer in the 2^nd^ decision assigns a prior probability (grey line) equal to the confidence in the first choice to positive signal values. Inasmuch as the response to the first task was correct, the true value of *s*_2_ for the 2^nd^ decision is positive (lower mid-panel). *s*_2_ being however small (as illustrated), it evokes by chance (due to internal noise) a negative internal response *r*_2_ that would (in the absence of prior information) lead to the erroneous conclusion that *s*_2_ was negative. Nevertheless, given the asymmetrical prior, the posterior distribution is still favouring the correct choice that *s*_2_ is positive. The lower left panel illustrates how a non-Bayesian observer (who does not have knowledge about the statistics of the internal noise, and only perceives point estimates) could make the same choices as the Bayesian observer by comparing the internal response *r*_1_ with a confidence criterion (the two vertical blue lines symmetrically placed about the decision criterion, dashed vertical line). By classifying decisions in two discrete levels, ‘confident’ vs. ‘uncertain’ or ‘non-confident’, and by shifting the 2^nd^ decision criterion only after ‘confident’ 1^st^ decision, the non-Bayesian ‘discrete’ observer can also increase the frequency of correct second choices. **B**. Relationship between confidence and internal sensory evidence for the Bayesian and non-Bayesian models, and for another model of a Bayesian observer that has a biased estimate of the internal variability, and can therefore display over or under confidence (relative to optimal). For the discrete and the biased Bayesian models the curve is computed by averaging MLE estimates of the parameters across subjects. Note that the discrete model (blue curve) does not actually compute probabilities, however it adjusts the criterion for the second decision. This criterion shift can be transformed into the equivalent confidence level of the ideal Bayesian observer.

**Table 1:** Results of the logistic analysis that measured the influence of the correctness of the first decision on the probability of choosing ‘right’ in the second decision (after accounting for the effect of stimulus values). The reported odds-ratio is between the odds of having the observer choosing ‘right’ (in the second decision) after a correct first decision and after a wrong one (given the same set of stimuli). The column *p*(correct) displays the mean and standard deviations of the proportion of correct responses in the second decision, computed across observers. The last two columns are 95% CI of the odds-ratio for each condition.

| Condition | Experiment | $p(\text{correct})$ | $\chi^2$ | $p$ | odds-ratio | 95%LCL | 95%UCL |
| --- | --- | --- | --- | --- | --- | --- | --- |
| O-O | correlated-pairs | 0.86(0.05) | 12.50 | 0.0004 | 6.21 | 3.09 | 12.47 |
| O-O | random-pairs | 0.88(0.05) | 1.23 | 0.27 | 3.47 | 0.43 | 27.97 |
| D-D | correlated-pairs | 0.88(0.06) | 14.16 | 0.0001 | 10.12 | 4.88 | 21.00 |
| D-D | random-pairs | 0.87(0.04) | 16.02 | <.0001 | 32.75 | 15.82 | 67.76 |
| D-O | correlated-pairs | 0.85(0.07) | 14.37 | 0.0001 | 9.68 | 4.59 | 20.42 |

### Computational models

Given the specific stimuli presented in the first decision and participant’s trial-by-trial responses, participant’s performance in the dual-decision paradigm can be compared with that of an ideal observer that accurately estimates the probability of being correct in the first decision and uses it as prior information for the second decision, according to the rules of Bayesian decision theory (see Figure 2A top). Assuming that the sensory noise is adequately described by a Gaussian distribution, updating prior expectations amounts to shifting the decision criterion for the second decision by the same distance observed between the first signal and criterion, expressed in standard deviations of sensory noise, *σ* (see Supplemental information for the full derivation). We have also compared human’s behavior with that of three other classes of models: (i) a biased Bayesian observer model, which also performs probabilistic computations to estimate its confidence, albeit using a biased estimate of the variability of its internal response (this allows the model to reproduce patterns of over or under confidence, which would correspond respectively to under or over estimation of the variability of the internal noise); (ii) a heuristic, or non-Bayesian observer characterized by a finite number of discrete confidence levels (this model implements the idea that observers may make perceptual decisions using only point estimates of the decision variable, rather than the full probability distribution, see Figure 2A bottom); and (iii) a model representing a sample-based approximation of the Bayesian model (where the posterior distribution is approximated by finite number of samples; see Supplemental information). These models made different predictions of the relationship between sensory evidence and confidence (see Figure 2B) and can be reliably discriminated in synthetic datasets (see Supplemental information, Figure S3). Compared to the optimal Bayesian model, the biased Bayesian model is characterized by one additional free parameter, a scaling factor that indicates by how much the participants over or under estimate the variability of their sensory noise. The non-Bayesian models instead are characterized by two additional free parameters for each additional discrete confidence level: the first parameter indicates the location of the confidence criterion (vertical blue lines, Figure 2A bottom), and the second parameter indicates by how much the criterion for the second decision is shifted following a ‘confident’ first decision (black dashed vertical line, in Figure 2A bottom-right). A summary of the estimated values of model parameters is reported in Supplemental information, Table S2.

### Model-free analyses

Figure 1B shows the proportion of correct responses as a function of discriminability (measured in units of internal noise) in the first and second decision (see Figure S1 for separate plots for each experiment and condition). As it can be seen, second decision performance (darker traces) is higher than first decision performance, especially for the most difficult trials (where the difference between the two stimuli in a pair was very small and first decision performance was close to chance). Since the more tilted or longer stimuli appeared more often in the right placeholder for the second decision (given that task difficulty was adjusted so as to yield average first-decision performance above chance), an increase in second-decision performance could be the result of a fixed bias (i.e., participants having chosen a ‘right’ response more frequently), without necessarily involving a trial-by-trial monitoring of uncertainty. To control for this possibility we performed a logistic analysis to measure the influence of the correctness of the first decision on the probability of choosing ‘right’ on the second decision (i.e. reporting that the signal was drawn from the positive semiaxis, that is *s*_2_ > 0). For each experiment (correlated- and random-pairs) and condition (O-O, D-D, and D-O) we fitted a multilevel (mixed-effects) logistic regression, using **R** [36] and the lme4 package [2], with the absolute difference between the two stimuli (in units of *σ*) and the accuracy of the first response as fixed effect predictors, and the participant as a grouping factor. We evaluated statistically the effect of the correctness of the first response with a likelihood ratio test between the fitted model and a reduced model where the effect of the first response was set to 0. This test was significant for all the experiments and conditions (see Table 1), with the exception of the O-O condition in the random-pairs experiment. In order to check whether a simpler fixed-bias model would really suffice to describe performance in this latter case, we performed a Monte Carlo simulation. For each trial we estimated the expected probability of a ‘right’ second response on the basis of the stimuli presented and the psychometric function fitted to the first decision responses. This results in a set of Bernoulli trials with different probabilities of success, which we simulated 10^5^ times in order to estimate, using the percentile method, a 99.5% confidence interval (corresponding to the Bonferroni corrected alpha level 0.005) on the expected proportion of second responses ‘right’ given the stimuli and the first response. The results revealed that after a wrong first response none of the participants responded ‘right’ more often than what would be expected given the stimuli: all the observed proportions were within the confidence intervals. Instead, after a correct response, the observed proportion of responses ‘right’ exceeded the confidence interval for 3 out of the 5 participants in this experiment. This result suggests that also in this condition, where the fixed bias hypothesis could not be rejected at the group level, the evidence favors the hypothesis that most participants have monitored the confidence in their first response on a trial-by-trial basis.

**Figure 3:**
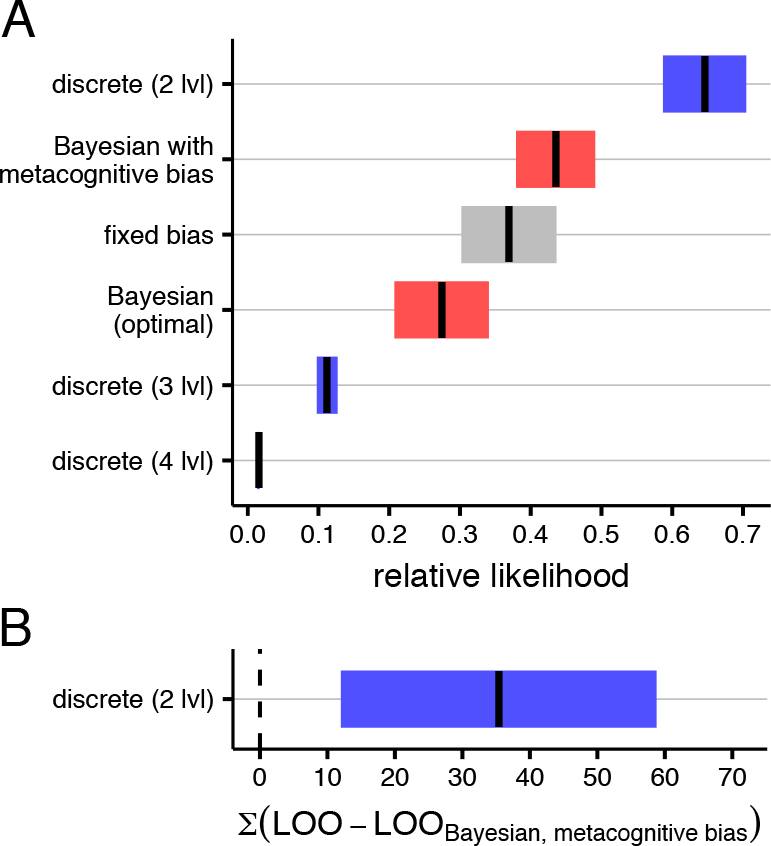
Model comparison. **A**.Relative likelihoods 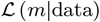, averaged across participants (see main text for details). Models depicted in red are probabilistic models which use full probabilistic distributions over the decision variable; in blue are models that only use point-estimates, but still evaluate the uncertainty heuristically on a trial-by-trial basis; the model in grey represents an observer that does not monitor uncertainty but simply adopts a fixed bias, corresponding to the belief of being more frequently correct than wrong in the first decision. Black lines represent mean likelihoods and error bars represent bootstrapped standard errors of the mean (SEM). **B**. Leave-one-out (LOO) cross-validated log-likelihood differences (discrete 2lvl minus Bayesian with a metacognitive bias) summed over subjects and task (the error bar is the bootstrapped standard error of the sum).

Overall, choice accuracies in the second decision revealed only small differences across task (see the also table 1). We investigated also whether our manipulation of the statistical relationship between the difficulties of first and second decisions did affected performance. We run a multilevel logistic regression, with the accuracy of the second decision as dependent variable, and the stimulus differences (absolute values) together with the experiment (random vs. correlated pairs) as predictors. This analysis did not reveal any significant difference in performance, *χ*^2^(1) = 0.55, *p* = 0.46, therefore data from the two experiments were collapsed in subsequent analyses. We used the same method to compare conditions where the two decisions were made in the same modality (O-O and D-D) with the condition where they were made on different modality (D-O; for this comparison only data from the correlated pairs experiment were used). Interestingly, we found that the difference in accuracy between these two conditions, although small (≈ 2% see table 1), resulted statistically significant *χ*^2^(1) = 8.24, *p* = 0.004. We will discuss the possible implications of this finding in the Discussion.

**Figure 4:**
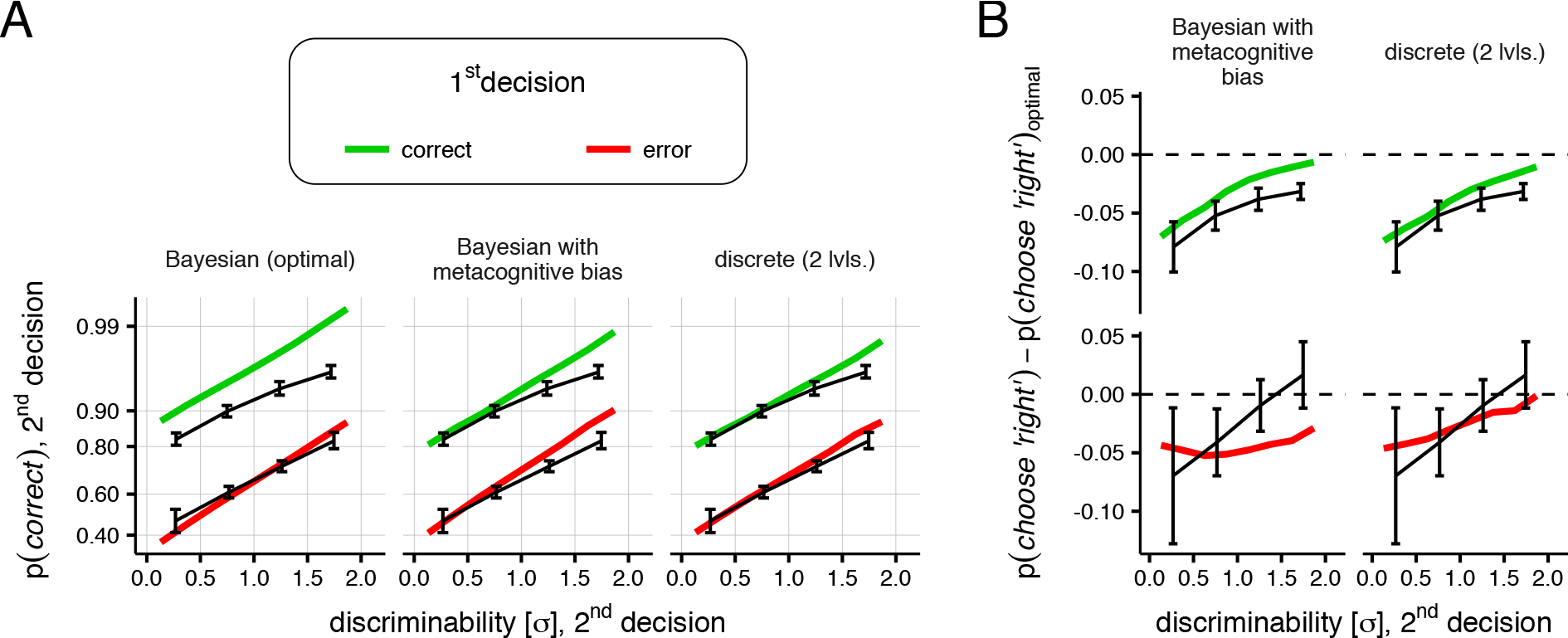
Comparison of model predictions and empirical data. **A**. The predicted (colored lines) and observed (black lines, with error bars representing ±1bootstrapped SEM) proportion of correct choices (i.e. reporting *s*_2_ > 0), split according to the correctness of the first decision and averaged across all observers and conditions, are plotted on a probit scale as a function of the discriminability of the second pair of stimuli (expressed in units of internal noise, *σ*). **B**. Differences in the proportion of choices ‘right’ of the biased Bayesial model (left panels), and of the discrete model with 2 levels of confidence (right panels). The upper panels demonstrate the under-confidence (relative to optimal), present in the data and reproduced by both models. The lower panels display choices after a wrong first decision, demonstrating the pattern of high confidence errors: after a wrong first response, subjects do not show the same pattern of under-confidence as after a correct first response. Instead, they tend to choose ‘right’ (and therefore commit an error) almost as often as the ideal, particularly for easy second decisions.

### Model comparison

We compared the models using the Akaike Information Criterion, AIC [1], to account for the different number of free parameter. For ease of interpretation, AIC scores were converted to a probability scale by taking the difference with respect to the AIC of the best fitting model (separately for each participant) and transforming the result into relative likelihoods of the model given the data, 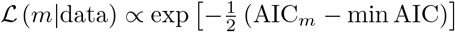 [8]. We also included into the model comparison a fixed-bias model, which corresponds to the hypothesis that observers did not monitor their own uncertainty on a trial-by-trial basis, but simply increased the frequency of responses ‘right’ in the second decision by shifting their decision criterion by a fixed amount. Although this hypothesis was rejected by model-free analyses, it is useful to see how it compares to the other models. We find that the model with the higher relative likelihood was the non-Bayesian observer with 2 discrete confidence levels (Figure 3A). In particular, the relative likelihood (averaged across tasks and conditions) of the non-Bayesian observer with 2 confidence levels was higher than the ideal Bayesian observer for 13 out of 14 observers. Most interestingly, it was also higher than the second best model (the Bayesian model with metacognitive bias) for 11 out of 14 observers. As an additional test we run a leave-one-out (LOO) cross-validation of the two best fitting models, and found again that the non-Bayesian model was on average better at predicting the actual behavior of the participants (Figure 3B).

Given the results of the model comparison, it is important to understand the differences between the predictions of the Bayesian and non-Bayesian models. Figure 4A displays on a probit scale the observed proportion of correct second decision choices as a function of the difficulty of the first decision, averaged across observers, tasks and conditions. The examination of this plot reveals several interesting features. First, the left panel in Figure 4A indicates that the main difference between the data and the prediction of the ideal Bayesian observer is that human observers make more frequent errors than the ideal after a correct first decision, which implies that they choose ‘right’ less frequently than ideal. Because in our task the bias favouring ‘right’ responses in the second decision is a function of the confidence in the first decision (see Supplemental information and Figure 2), this pattern denotes *under confidence*, i.e. human observers consistently underestimate their probability of being correct in the first decision. The finding of a global under-confidence bias allows us to reject the alternative models based on a sampling scheme, in which probability distributions are approximated by a finite number of samples. We find that sampling-based approximations of the ideal Bayesian model cannot adequately describe the data, at least not without a number of additional assumptions, because for a small number of samples they produce a marked overconfidence bias (see Supplemental information), opposite to the one found in the data. Note that as the number of samples increases the sampling models converge to the ideal Bayesian model. The biased Bayesian observer instead can account for this pattern, by assuming that participants use a biased estimate of their sensory noise (figure 4A, middle panel); maximum likelihood estimation of this parameter indicates that the under confidence bias in the data could be explained by assuming that they over-estimate their sensory noise approximately by a factor of 3.

The biased Bayesian observer is less confident than the ideal also after wrong first decisions, leading to less frequent errors in the second decision^1^ This implies that after an error, human observers respond more frequently ‘right’ than predicted by the biased Bayesian model, indicating that they make more frequent high-confidence errors (or alternatively that they are on average more confident when they make a wrong choice). This can be appreciated better in Figure 4B, where the predicted and observed rates of ‘right’ second decision choices are represented as differences from the ideal. The upper panels show the under-confidence pattern (after correct first decisions). This is predicted by both the biased-Bayesian and the alternative, non-Bayesian discrete model. The lower panels of Figure 4B show that the under-confidence is not systematically present after a wrong first decision; instead, observers deviate from the predictions of the biased Bayesian observer and respond more frequently ‘right’; in particular they chose ‘right’ as frequently as the ideal for trials where the intensity of the second signal was high. This implies that they were less sensitive than the biased Bayesian observer to the evidence provided by the second pair of stimuli when their first decision was wrong.

Only the discrete model predicts rates of ‘right’ responses that are closer to the observed ones, simultaneously showing underconfidence after correct first decisions, and more high-confidence errors. According to the discrete model, high-confidence errors are to be expected whenever the internal signal not only has opposite sign with respect to the physical stimulus (due to random noise fluctuations), but is also large enough to exceed the confidence threshold (see Figure 2) and consequently elicit a ‘confident’ wrong decision. The relationship between the internal signal, or sensory evidence, and the confidence predicted by the ideal Bayesian observer and the two best fitting models is plotted in Figure 2B: high confidence errors occur whenever the signal upon which the wrong decision is based falls in the high-confidence region of the discrete model (blue, stepwise curve). With our behavioral task we cannot measure the trial-by-trial internal signals, however we can compute their expected value as the mean of their probability distribution conditioned on the stimulus and the choice (this amounts to taking the mean of a truncated Gaussian distribution centered on the true value of the stimulus with upper or lower boundary at 0, depending on whether the observer decided ‘left’ or ‘right’, respectively) and compare it to the individual confidence thresholds estimates. This analysis revealed that even for wrong first decisions, the internal signal was expected to exceed the confidence criterion in ≈ 42% of the trials (and in ≈ 77% of correct first decisions), supporting the idea that a substantial proportion of high confidence errors is to be expected given the stimuli and the observed pattern of choices.

## Discussion

We developed a novel approach where in a sequence of two dependent perceptual decisions humans could improve their second decision performance by taking advantage of the fact that the statistics of the stimuli presented for this second decision depended on the correctness of their first decision. This experimental protocol can be regarded as a laboratory proxy of more complex environments in which confidence is used to guide behavior in cases where a current decision depends on the unknown outcome of a previous one. We developed a normative Bayesian observer model for this task, i.e., an ideal observer who performs Bayesian inference to estimate the posterior probability of being correct in the first decision (the confidence), and uses it optimally (i.e., maximizes the probability of a correct second decision) as a prior probability distribution for making the second decision. By comparing participants’ performance with the predictions of the Bayesian model, we were able to test the hypothesis that human observers are able to evaluate probabilistically their own uncertainty and make meta-perceptual judgments according to a Bayesian-probabilistic strategy.

The present results revealed clearly that the participants engaged in this task exhibited systematic deviations from the predictions of the normative Bayesian model. While they were clearly able to take trial-by-trial uncertainty into account (a simple fixed bias model was not sufficient to account for the observed behavior: see Results, Model-free analyses), their pattern of second decisions revealed the presence of a marked, global miscalibration of confidence, more frequently in the form of under-confidence, as indexed by lower rates of responses ‘right’ in the second decision with respect to ideal. We developed an alternative ‘biased’ Bayesian model, designed to reproduce global miscalibration of confidence such as the observed under-confidence by assuming that observers use a Bayesian-probabilistic strategy but may have a biased estimate of the variability of the internal signals. This model is in line with recent work showing that transcranial magnetic stimulation (TMS) on the visual cortex can degrade performance in a visual discrimination task without affecting comparative judgments of uncertainty made over multiple decisions [34]. Peters and colleagues [34] explained the results by assuming that human observers rely on a learned statistical model of their own perceptual system that may become invalid under specific conditions, such as the external disturbances elicited by means of the TMS. In that case subjects are metacognitively blind to the additional noise introduced by the TMS and therefore under-estimate their internal variability. In our case, the biased Bayesian observer model can reproduce the under-confidence bias by assuming that, on average, participants over-estimate their internal noise approximately by a factor of 3.

Although the biased Bayesian outperformed the ideal, unbiased Bayesian model in predicting the actual behavior, we found additional discrepancies between predictions and empirical data that question the biased-Bayesian’s descriptive accuracy. Specifically, we found that on average the biased Bayesian model predicts less frequent (with respect to behavior) ‘right’ choices after a wrong first decision, see Figure 4, indicating that human observers made more frequent high confidence errors. Overall, our results show that (i) human observers show less confidence than the ideal Bayesian observer after a correct first decision, but at the same time (ii) after a first wrong decision are more confident than the alternative Bayesian model augmented with a bias parameter to account for the underconfidence (relative to ideal). Taken together, these findings indicate that the relationship between the internal signals (the sensory evidence) and the confidence does not have the shape that should be expected if subjective confidence were computed as by a Bayesian observer, namely as the posterior probability of being correct.

The pattern found in our data could be accounted for by our non-Bayesian model, in particular the simple model with only two discrete confidence levels^2^ From a psychological point of view, the non-Bayesian class of models posits that confidence is discretized in a number of distinct levels, as also suggested by previous work [44]. For example, in the one-criterion variant of our model there would be two discrete confidence states: confident - i.e more likely that the response was correct - vs. non-confident - i.e., equally likely that the response was correct or wrong (see Figure 2). This one confidence criterion variant of our model provided the best and most parsimonious description of the empirical data (see Results, Model comparison). It is important to note that because this model does not assume any knowledge about the variability of the internal decision variable, these confidence levels are defined only on an ordinal scale and do not convey a precise numerical information about the probabilities involved.

Although our heuristic class of models may seem too simplistic to represents an accurate algorithmic description of the mechanisms underlying subjective confidence valuation, we argue that the specific way in which the relationship between sensory evidence and confidence is realized in this class of models - as a discrete step function - must capture some aspects of the actual mechanism implemented by the brain. More specifically, we propose that confidence grows with sensory evidence according to a piecewise function: more slowly than ideal for a certain range of values, or sub-domains, of sensory evidence (e.g., for values close to the decision boundary 0, contributing to produce the global under confidence bias), but at the same time it also grows faster than ideal in other sub-domains, resulting in more frequent high confidence wrong decisions. This proposal implies that subjective confidence does not match the Bayesian posterior probability of the ideal Bayesian observer, but instead is based on different computational processes that may require only point estimates, as opposed to full probability distribution. This pattern where subjective probabilities both overestimate and underestimate the objective probability in distinct sub-domains of the same task resemble the distortions of subjective probability that have been found also in non-perceptual task, such as those involving decisions from experience [45].

Given that participants’ performance was slightly lower in the D-O condition (see Results, Model-free analyses), where first and second decisions were made in different modalities, it is reasonable to ask whether differences of the observed data from the predictions of the Bayesian models were mostly driven by this specific condition. Interestingly, looking at the model likelihoods separately for each condition, see Figure S4 it becomes evident that this was not the case as the D-O condition was instead the only one where the likelihood of the biased-Bayesian resulted slightly higher than that of the non-Bayesian model with two confidence level. Taken together, both the pattern of model likelihoods, and the slightly lower performance, suggest that participants may have used a slightly different strategy in this condition. While our current data do not allow to pinpoint exactly what caused these different outcomes, they do suggest that claims of confidence as a ‘common currency’ between different perceptual judgments [12,13] should be re-evaluated using a more diverse set of experimental protocols.

Taken together, our behavioral and modelling results suggest that humans can use sensory evidence to perform probabilistic judgments, but ultimately cannot assign numerically precise subjective probabilities to perceptual interpretations. These comparative probability judgments, not linked to precise numerical values, are the essence of qualitative probability reasoning [39], a weakened but more pragmatic and intuitive counterpoint of classical probability theory. The notion of qualitative probability lies at the foundations of the notion of subjective (or Bayesian) probability since its early formulation [11,39]. Much of the early work on the theory of Bayesian probability has been dedicated to identifying the conditions that allow the departure from the qualitative probability toward the quantitative (numerically precise) probability, defined according to the classical Kolmogorov axioms [24]. Several propositions have been put forward, but all of them ultimately assume that the decision-maker’s knowledge allows the partition the probability space associated with the event space into a uniform and arbitrarily large collection of disjoint states or events [39]. While this assumption can be used to provide a formal framework for exact reasoning under uncertainty, it may be too fine-grained for a realistic, biological decision-maker. Indeed, in the real world assigning exact numerical probabilities is often difficult or impossible, and the ability to compare the likelihood of two events without having to provide exact probabilities may be sufficient. Our results support this idea, as the performance of the heuristic observer with two discrete confidence levels, despite its relative simplicity, shows a marked increase in the probability of correct second decisions relatively to a simpler model which does not consider any prior information (the expected proportion of second correct choices in the absence of any criterion shift was ≈80%), while the ensuing performance benefit obtained by the Bayesian model may not be enough to justify its increased computational complexity [25] and costs. Indeed, it is known that efficient adaptive decisions can be taken also by means of simple heuristics, which may outperform more rigorous Bayesian strategies once the costs of information acquisition and processing are taken into account [18].

Our findings suggest that subjective confidence does not match the Bayesian posterior probability of being correct, even after taking into account the possibility that observers have a biased estimate of the internal noise or approximate the subjective probabilities with a finite set of samples. These results have important implications for the current investigation of the neural substrate of metacognition, given that many of the relevant studies assume, more or less explicitly, that subjective confidence corresponds to the Bayesian posterior probability (e.g., [23,30]). Although Bayesian decision theory provides a general and logic way of processing information, the evidence supporting the hypothesis that it is algorithmically implemented in the brain is debated [26,37]. We argue that to achieve real progress in the understanding of human decision-making and metacognition, it is fundamental that researchers consider also alternative hypotheses and models. Moreover, the view that Bayesian decision theory is always a normative description of rational decision making (sometimes referred to as “Bayesianism”) has been criticized also on purely theoretical grounds [3]: while Bayesian decision theory is demonstrably optimal in the context of “small” worlds [4,39] where all relevant alternatives, consequences and probabilities can be meaningfully estimated and assigned, it is unclear to what extent it can be considered rational in the context of “large” worlds, where not all the alternatives, consequences and probabilities are known. While simple perceptual inference problems can, under certain conditions, be effectively considered “small world” problems, there is no real reason for why this should be the case of perception in general. This line of reasoning thus undermines the proposal that subjective confidence should necessarily be operationalized as the Bayesian posterior probability, and motivates the need of empirical studies and novel approaches to test the descriptive accuracy, as well as the ecological rationality [9] of applying Bayesian decision theory as a *process* model [27] to explain the behavior of imperfect, biological decision makers. In the present study we have not only found further evidence that brain processes subtending meta-perception do not conform to the ideal benchmark represented by the Bayesian observer, but also provided a novel, general experimental protocol that we believe will be a valuable tool for future investigations of confidence and metacognition. The availability of a rich set of new protocols for assessing cognitive functions is also important for translating advances in neuroscience and cognitive science into concrete, theory-driven, clinical applications [21].

## Ackowledgments

This work was supported by Grant ANR-12-BSH2-0005 from the French National Research Agency to A. Gorea.

## Material and methods

Participants (5 for the random-pairs experiment, and 9 for the correlated-pairs experiment) performed a sequence of two decisions on each trial, with the statistics of the second decision stimuli being conditioned on the correctness of the first decision, with this dependence made explicit to the observers. All experiments were performed in accordance with French regulations and the requirements of the Helsinki Convention. The protocol of the experiments was approved by the Paris Descartes University Ethics Committee for Non-invasive Research (CERES).

We describe here the general structure of the protocol, with the details of the implementation and of the analysis being provided in the Supplemental information. At the beginning of each trial, two stimuli were presented in two placeholders on the left and right of the fixation point. The two stimuli differed along one physical dimension [orientation of the Gabor-patches - spatial frequency 1.5 cycles/dva (degrees of visual angle), standard deviation of the envelope 0.7 dva, contrast 25%, - or duration - Gaussian blobs with a pick intensity of ≈ 25.3 cd/m^2^ and a standard deviation of 0.65 dva]. The participant was required to indicate which of the two stimuli was characterized by a higher value along the given dimension by pressing the left/right arrow keys. The difference between the two stimuli was uniformly distributed within 2 JNDs (just noticeable differences), measured in preliminary sessions (see below). 400 ms after providing the first response, a second pair of stimuli was presented and the participant was again asked to indicate which of the two has a higher value. The difference in value between the stimuli in the second pair was also randomly sampled from a uniform distribution. However the location of the higher-value stimulus depended this time on the correctness of the first response: if the first response was correct, the higher-value stimulus was presented on the right, and on the left otherwise. Participants were informed about this rule, and were asked to use it in order to achieve the best possible second decision accuracy. Before starting the experimental trials, participants were explained the rule and were familiarized with a version of the task where the difference between the two stimuli to be compared could go up to very high values (up to 45° in the orientation task, and up to 1 second in the duration task, uniformly distributed). The large differences in the practice session were intended to make the rule clear and unambiguous for all participants. In one first experiment discrimination difficulties in the first and second decisions were drawn independently (random-pairs). In a second experiment we biased the probability of association between discrimination difficulties in the first and second decision (correlated-pairs, see Supplemental information, Figure S5). Specifically, when the difference in intensity in the first pair was less than 1 JND, there was a 0.7 probability that the difference in the second pair would be larger than 1 JND, and vice versa. This was intended to encourage participants to make use of the rule. The random-pairs experiment was tested in two conditions, run in different sessions on different days (order balanced), where the two decisions involved both an orientation discrimination (O-O) or a discrimination of duration (D-D). Each of these sessions comprised 500 trials. The correlated-pairs experiment was declined in three different conditions, O-O, D-D and D-O (where the two decisions involved a duration discrimination followed by an orientation discrimination). Each of the three sessions comprised 300 trials, each consisting of two consecutive perceptual decisions. The different number of total trials in the correlated- and random-pairs experiments was designed so that they resulted in similar number of easy (difficult) decisions followed by difficult (easy) decisions. Each testing session was divided in 10 blocks of trials. At the end of each block participants were given a feedback about the overall accuracy of their second decisions in that block. Additionally, to help participants keep track of their performance, starting with the end of the second block they were also informed on whether their accuracy had increased or decreased with respect to the previous block.

## Analysis

For each participant we estimated the standard deviation of the internal noise, *σ*, by fitting on the proportion of ‘right’ choices in the first decision four different psychometric models that made different assumptions about whether participants were biased or made frequent attention lapses (i.e., stimulus independent errors), and by combining the 4 estimates of the standard deviation by weighting them according to the Akaike weight of each model [8] (see Supplemental information for details). Next, we used the estimated *σ* to transform the values of the stimulus from raw units (e.g. degrees and seconds) to units of internal noise. Finally, we fitted the non-Bayesian models using maximum likelihood estimation, and compared the predictions of Bayesian and non-Bayesian models with the observed pattern of second decisions (all models make similar predictions for the first decision). Because these models differ in the number of free parameters, in order to prevent overfitting we compared the models on the basis of the Akaike Information Criterion [1]. Additionally we performed a leave-one-out cross-validation of the two best-fitting models. All analyses were performed in the open-source software **R** [36]; the data and the code of the analysis are available upon request. The mathematical details of the computational models are provided in Supplemental information.

## Supplemental information

### Material and methods

#### Apparatus

Participants sat in a quiet, dimly lit room, with the head positioned on a chin rest at a distance of 60 cm from the display screen, a gamma-linearized Mitsubishi Diamond Plus 230SB CRT monitor (screen resolution 1600×1200, vertical refresh rate 85 Hz). Stimuli were generated by a computer running Matlab (Mathworks) with the Psychophysics Toolbox [7,33].

#### Stimuli and procedure

All experiments were run with the same visual display, consisting of a central fixation point and four placeholders, continuously visible on a uniform gray background (luminance ≈ 13.6 cd/m^2^). The four placeholders were circles measuring 2.8 dva (degrees of visual angle) in diameter, whose centers were placed at 1.8 dva from the fixation point. Two placeholders were grey (≈ 15.5 cd/m^2^), and were placed above the horizontal mid-line; the other two were placed below the mid-line and were colored in red the one on the left, and in green the one on the right (their luminance was matched with the grey placeholders, ≈ 15.5 cd/m^2^).

In the orientation (O-O) task the stimuli were two Gabor gratings (sinusoidal luminance modulation presented within a Gaussian contrast envelope) of different orientations, presented for 200 ms. The spatial frequency of the Gabors was set at 1.5 cycles/dva, the phase was drawn randomly, and the standard deviation of the Gaussian envelope was 0.7 dva. The Gabor displayed in the left placeholder was always tilted to the left, and the one appearing in the right placeholder was always tilted to the right. The task of the participants was to indicate which Gabor was more tilted from the vertical; the less tilted of the two Gabors was always tilted by 15°; the minimum difference was 0.1°.

In the duration (D-D) task the two stimuli consisted of white Gaussian blobs (standard deviation 0.65 dva), presented sequentially in the two placeholders (left/right). The order of presentation (left/right stimulus first) was balanced with respect to the longer/shorter duration of presentation. Participants were asked to indicate the location (left/right) of the longer duration blob. The shorter duration was always set to 600 ms, and the difference between shorter and longer durations was discretized in bins determined by the vertical refresh of the monitor (≈ 12 ms). The minimum duration difference was one single monitor refresh interval.

#### Pre-test JND measurement

Before the orientation and duration task, we measured individual JNDs using a weighted up-down staircase procedure [22]. The purpose of this pre-test was to quickly obtain a measure of the JND in order to adapt the range of stimuli in the main experiment to individual sensitivities. The staircase procedure continued until 30 reversals were counted. The initial step size (the size of the decrease/increase of the difference between the two stimuli) was 2° in the orientation task and 4 refresh intervals in the duration task (≈ 50ms), and was diminished to 0.5° and 1 refresh after the second reversal. Stimuli in these pre-test measurements were presented only in the top placeholders.

#### Participants

5 subjects (2 female; mean age 30.8, standard deviation 3.1) participated in both conditions of the random-pairs experiment (D-D, and O-O; see Main text, Material and methods). 9 participants (4 female, 2 authors; mean age 33.9, standard deviation 9.6) participated in the 3 conditions of the correlated-pairs experiment (D-D, O-O, D-O). All participants (except the author) were naíve to the specific purpose of the experiment. All conditions were performed in separate session on different days. The order of the D-D and O-O conditions was counterbalanced across subjects, while the D-O condition was always performed in the last session. All participants had normal or corrected-to-normal vision and gave their informed consent to perform the experiments.

### Computational models

#### Bayesian observer

To model the performance in our task, we considered that the observer to make a decision estimates the difference in the intensity between the left and right stimuli, 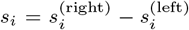, where the subscript *i* = 1, 2 indicates the decision at hand (first and second) and 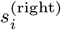 and 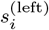 denotes the stimuli (unsigned deviations from vertical of the two Gabor gratings in the orientation task, or durations of the two blobs in the duration task). Hence the task amounts to deciding whether si is greater or less than 0. We assumed that the observer has only access to a corrupted version of *s*_*i*_, *r*_*i*_ = *s*_*i*_ + *η*, where *η* is Gaussian noise with variance *σ*^2^. The ideal Bayesian observer has full knowledge of the statistics of the internal noise, and to make a decision computes a posterior probability over the variable s. Since the internal noise is assumed to be Gaussian, the likelihood function giving the probability of observing *r*_*i*_ given *s*_*i*_ is 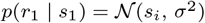, where 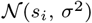 is a normal distribution with mean si and variance *σ*^2^.

In first decision, the prior probability of *s*_1_ is uniform with the range (−*R*, *R*), that is *p*(*s*_1_) = 1/(2*R*) if – *R* ≤ *s* ≤ *R* and 0 otherwise. Combining the prior and the likelihood function, the unconditioned probability of observing r1 can be expressed as:

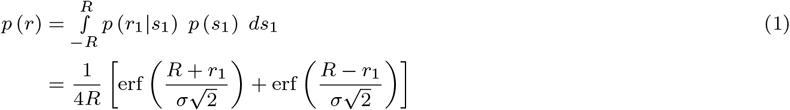

The posterior probability of *s*_1_ after having observed *r*_1_ is obtained applying Bayes rule:

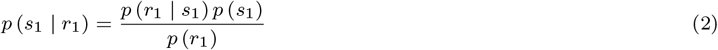

Finally, the decision variable 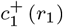, corresponding to the probability that the *s*_1_ was greater than 0 is given by

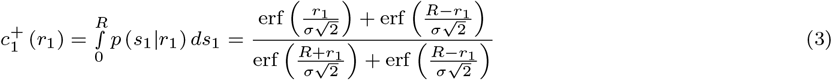

When 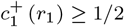 the observer chooses ‘right’ (i.e. he decides to report that *s*_1_0 was positive, an outcome hereafter indicated with the notation 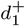) otherwise he choses ‘left’ (i.e. he reports that *s*_1_0 was negative, notated as 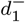). The posterior probability of being correct in the first decision, that is the confidence of the ideal observer, is given by 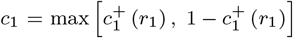. The ideal observer would use the probability *c*_1_ to adjust prior expectations for the second decision, specifically by assigning a prior probability equal to *c*_1_ to the possibility that *s*_2_ will be drawn from the positive interval:

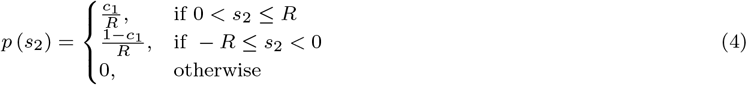

By applying the same calculation as above with the updated prior *p*(*s*_2_), one obtains the decision variable 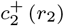 for the second decision:

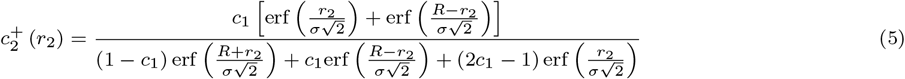

This equation reduces to the one for the first decision variable when *c*_1_ = 1/2, as it should. The range *R* on which *s*_1_ and *s*_2_ takes values is immaterial. It is possible to simplify Eqs (3) and (5) by taking the limit *R* → ∞. In this limit one obtains:

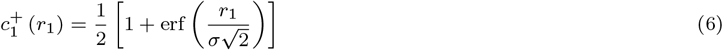

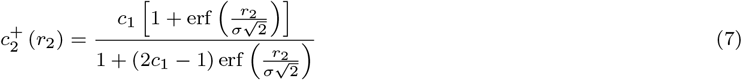

The decision rules described by these equations amount to comparing the internal signal *r*_*i*_ to a criterion *θ*, and decide accordingly (i.e., if *r*_*i*_ ≥ θ, chose 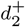, otherwise chose 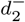). The criterion for the first decision, *θ*_1_, can be expressed as:

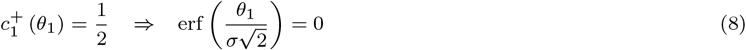

which is satisfied for *θ*_1_ = 0. Similarly, the criterion for the second decision, *θ*_2_, can be expressed as:

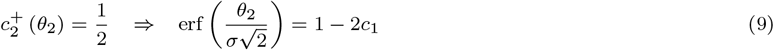

which indicates that *θ*_2_ is a function of *c*_1_ (see Main text, Figure 2B).

The likelihood of the Bayesian observer choosing 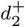 (after having choosen 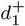 in the first decision) can be expressed as:

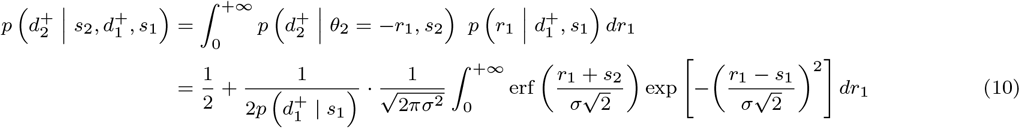

When the observer has chosen 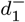 in the first decision instead the likelihood takes the form:

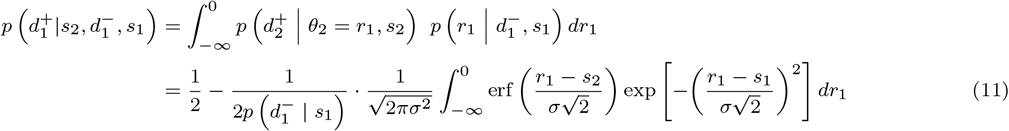

The integrals in the two equations 10 and 11 do not have closed form solutions, so they were evaluated numerically using the adaptive quadrature algorithm as implemented in the function **`integrate`**() in **R** [36].

#### *Biased*-Bayesian observer

Similarly to the ideal Bayesian observer presented in the previous section, the biased-Bayesian observer makes decisions based on an internal signal corrupted by Gaussian noise with variance *σ*^2^. However he does not know exactly the value of *σ* so when asked to assess his confidence he uses 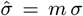 (with *m* > 0). Therefore, after having chosen 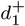 in the first decision, his confidence can be expressed as:

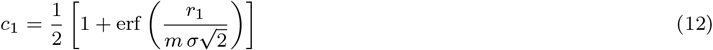

The adjusted criterion for the second decision is given by the same relationship that was found for the Bayesian observer

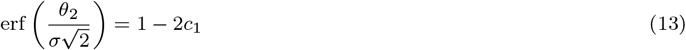

However in this case solving for *θ*_2_ gives

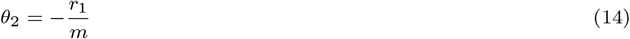

This means that when *m* > 1 the internal noise *σ* is overestimated 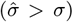 and the criterion is shifted away from zero less than in the optimal Bayesian model, leading to an under-confidence bias (i.e. in the second decisions the observer choses in agreement with the belief that the first decision was correct more frequently than what expected if performance was optimal). Conversely the criterion is shifted more than ideal when *σ* is underestimated (*m* < 1), leading to an opposite over-confidence bias.

#### Non-Bayesian observer

As an alternative to the optimal Bayesian model we considered a class of models that do not assume any knowledge about the nature of the internal stochastic process linking the stimulus *s*_*i*_ with the internal observation *r*_*i*_. These non-Bayesian models perform similarly to the Bayesian model for the first decision, that is when *r*_1_ ≥ 0 they chose 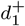, and 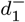 otherwise. However, they cannot estimate a full probability distribution over the values of *s*_*i*_, and therefore can assess confidence only by comparing the internal response *r*_*i*_ (which can be described as a point-estimate) to a set of one or more fixed criteria (Main text, Figure 2). In the case of a single confidence criterion, the non-Bayesian observer is confident in the response when the internal signal exceed the confidence criterion, and uncertain otherwise. When confident about the first response, he shifts the decision criterion for the second decision by a fixed amount (thereby increasing the probability of choosing 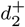). If only one criterion is used, then the model has 2 discrete confidence levels (e.g., uncertain vs confident). In such a model, if the confidence criterion is *w*_1_, the probability of the observer being confident about his first decision, after having responded 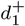, can be expressed as:

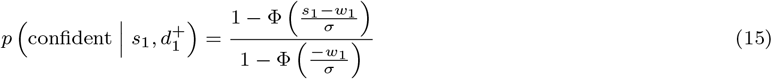

Note that this probability do not denote the confidence of the observer about his choice, which instead is assumed here to be a discrete binary state. Applying the law of total probability, the probability of the observer reporting 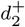 in the second decision can be expressed as:

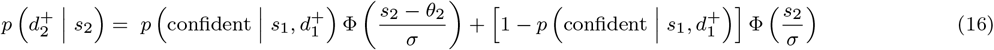

where *θ*_2_ is the shift in criterion for the second decision applied by the observer when he is confident in his first decision. *w*_1_ and *θ*_1_ are free parameters that we fit to the data by maximum likelihood estimation. It is straightforward to extend the model in order to have more than two confidence levels. In our analysis we considered models with 2, 3, 4 discrete levels of confidence, which had 2,4 and 6 free parameters, respectively. The parameters *w*_1_, *w*_2_, *w*_3_ and *θ*_2_, *θ*_3_, *θ*_4_ were constrained so that 0 ≤ *w*_1_ ≤ *w*_2_ ≤ *w*_3_ and 0 ≥ *θ*_2_ ≥ *θ*_3_ ≥ *θ*_4_.

#### Model fitting

In the first decision *s*_1_ is uniformly chosen from an interval centered around 0 (i.e., *s*_1_ will be above 0 with probability 1/2), and there is no prior information about the sign of *s*_1_. Therefore, for all the models presented in the previous sections, the probability 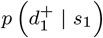 of the observer choosing 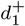 can be expressed as:

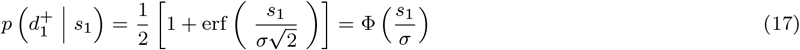

where Φ is the cumulative distribution function of the standard normal distribution. We fitted this function by maximum likelihood to estimate 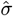, for each participant and task (duration and orientation), taking into account only the first decisions. We estimated also other psychometric models, that extended this simple model to account for the possibility that the observer were biased, 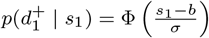 (where *b* indicate the bias), or made stimulus-independent errors (e.g. attention lapses) with nonzero probability, 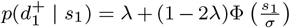 (where λ is the lapse probability); or both 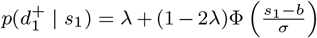. To avoid committing to any of these models, for each participant we averaged the estimates of 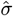 according to the Akaike weights [8] of each psychometric model, and used the model-averaged estimate for transforming the stimuli from physical units to units of internal noise. The same was done for the estimates of the bias term, 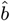, which was then subtracted from the stimuli value to take into account subjective decision biases in the computation of models’ likelihoods. The remaining free parameters for the biased-Bayesian and the non-Bayesian observer were estimated by numerically maximizing the (log) likelihood of the data (the observed patterns of second decisions). Maximum likelihood fits were obtained via the BOBYQA algorithm [35] as implemented in the **`optimx`** package [31,32] in **R**[36].

### Sampling-based approximation of Bayesian observer

An interesting alternative to the models presented in the previous sections is represented by models where the observer does not have access to the full probability distribution of his internal signals, but bases his decision on a limited number of samples. In these models the posterior probability that the choice is correct (the confidence) is approximated on the basis of a fixed number *n* of samples *x*_1_, *x*_2_, …, *x*_*n*_ drawn from the posterior distribution *p* (*s*_*i*_ | *r*_*i*_). The performance of these sampling-based models will approach the optimal Bayesian model as *n* ⟶ ∞, however they are expected to display systematic biases and deviations from the optimal model for small number of samples [38]. Here we show, and confirm by simulation, that the sampling-based approximation of the Bayesian observer will display a systematic over-confidence bias, that is in the opposite direction with respect to what we found for most of our subjects, and is thus inconsistent with our behavioral results.

### Sampling bias and over-confidence

It has been demonstrated that when a probability *p* is estimated from a small sample as the empirical frequency of successes *k* out of *n* random trials, *p̂* = *k/n*, the probability of overestimation, that is when *p̂* > *p*, or underestimation, *p̂* < *p*, depends in a complex way on both the probability *p* and the sample size *n* [41]. This is however for estimating a single fixed probability *p*. What would be instead the expected bias, over many repeated estimations, when the probability *p* varies randomly within a given range? In our experiments confidence is the posterior probability that a binary choice is correct, and as such it varies from complete uncertainty, *p* = 0.5, to complete certainty, *p* =1, hence *p* ∊ [0.5,1]. We show here that when a set of probabilities *p*_1_,…,*p*_*m*_ uniformly distributed in the interval [0.5, 1] is estimated using a limited number of samples *n*, the predominant bias is one of over-estimation.

For a given *n* and *p* the probabilities of over- and under-estimation can be expressed as:

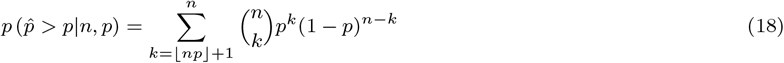

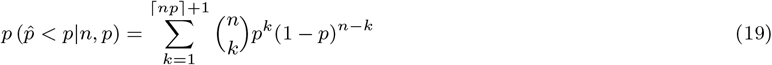

Where ⌈·⌉ and ⌊·⌋ are the ceiling and the floor operators, i.e. functions that map a number to the smallest following integer or the largest previous integer, respectively. Following Shteingart and Loewenstein [41] we consider the difference between these two, denoted as *probability estimation bias*, which takes the form:

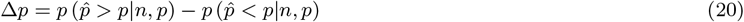

When Δ*p* is positive, it indicates that the probability *p* is more likely to be overestimated than underestimated, and viceversa for negative values. Assuming that all values of *p* in the interval are equally likely, the expected bias can be computed by integrating Δ*p* over the range of *p* (that is [0.5, 1]):

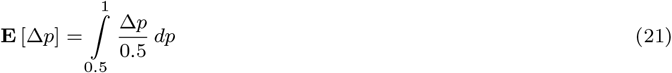

A positive value of the expected probability estimation bias (that is **E** [Δ*p*] > 0) indicates that, on average, the probabilities in this interval are more frequently overestimated rather than underestimated. This integral can be evaluated numerically, and in Figure S2A we plotted the expected probability estimation bias as a function of the number of samples, for two different ranges When p varies within the range of confidence, [0.5, 1], the value of the probability estimation bias is always positive, although modulated by the number of samples, indicating that in the range [0.5, 1] overestimation is more likely than underestimation.

### Fixed-*n* Bayesian sampler

Here we confirm by simulation the intuition presented in the previous paragraph. We consider a fixed-*n* policy, where the observer draws a fixed number of samples for each decision. Although alternative decision policies are possible (such as an accumulator policy, where the decision is taken after a minimum number of samples is accumulated in favor of one of the options), these have been shown elsewhere to result in very similar performances as the fixed-*n* policy [43].

We start by providing the mathematical details of the model. Similarly to the previous cases, we assume that the observer has only access to *r*_1_, a corrupted version of the stimulus *s*, *r*_1_ = *S*_1_ + *η*, where *η* is Gaussian noise with variance *σ*^2^. In the first decision the prior is flat and, taking the limit of the stimuli range *R* ⟶ ∞, the posterior distribution *p* (*s*|*r*) results in a Gaussian distribution centered on the internal observation *r*_1_. The probability that a sample from this distribution is above 0 (the criterion for the first decision) can be computed as:

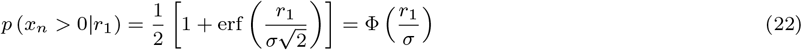

The conditional probability that the observer chooses 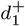 given *r*_1_ is obtained by summing the probability of all the set of samples with at least 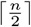 samples above 0 and can be expressed as:

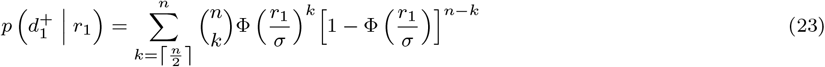

where *p* (*x*_*i*_ > 0|*r*) is the probability that a single sample *x*_*i*_ is above 0. In other words the observer choses 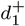 when the majority of samples is above 0. The observer’s confidence in his decision, 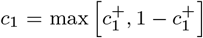, is given by the proportion of samples in favor of the choice made, which can be expressed as:

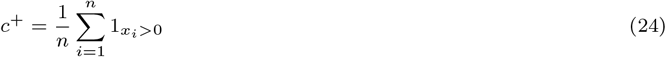

where 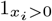 denotes the indicator function (1_*x*>0_ = 1 if *x*> 0, and 0 otherwise). The probability that the observer chooses 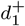 given *s*_1_ can be calculated by integrating the probability 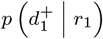 over all possible values of *r*_1_:

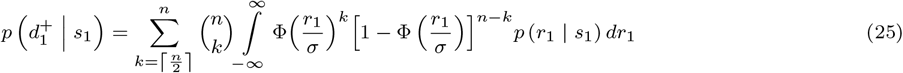

In the second decision the prior probability of the stimulus *s*_2_ is different for stimuli above or below 0. From the point of view of the observer, the prior probability that the stimulus for the second decision *s*_2_ is above 0 corresponds to the confidence *c*_1_ that the first decision was correct and, conversely, *p* (*s*_2_ < 0) = 1 − *c*_1_. Updating the prior probability for the second decision amounts to a shift in the decision criterion, as demonstrated for the full Bayesian model. The shift in criterion for the second decision *θ*_2_ is a function of the confidence in the sampling model as in the full Bayesian model

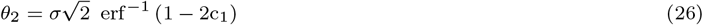

However, and differently from the full Bayesian model, the confidence is limited to a finite set of values determined by the number of samples *n*. The number of possible confidence levels is 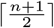, and each of these corresponds to a value of *θ*. If the observer’s first decision was 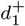, then the probability of the confidence level *c*_*j*_

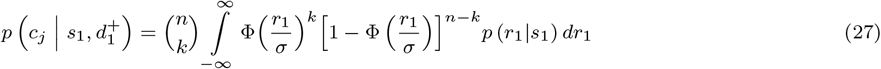

where 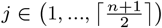 is an index linked to the number of samples above the criterion, *k*, according to 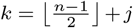. A level of confidence *c*_*j*_ would result in a shift in decision criterion *θ*_*j*_, calculated according to Eq (26). The probability of choosing (+) in the second decision is

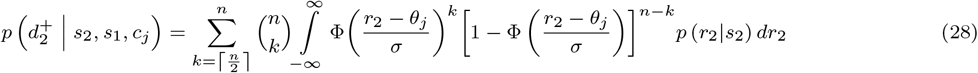

Taking everything together, the probability of the observer choosing 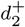 given *s*_2_, *s*_1_ and the first decision can be expressed as:

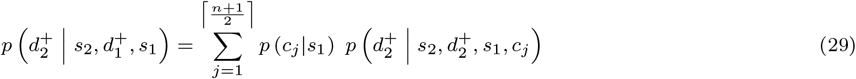

### Simulation

We simulated the model for values of *n* ranging from 2 to 9. In order to compare the model with the full Bayesian observer, the value of *σ* for each *n* are adjusted so as to obtain the same proportion of correct responses in the first decision. For each value of *n* and for 5000 iterations we: (1) generated a random set of stimuli for 500 trials; (2) simulated the Bayesian model on those trials; (3) estimated *σ* for the sampling models based on the set of first responses produced by the Bayesian model (this was done using maximum likelihood estimation and equation 25); (4) simulated the sampling model. The average values of *σ* obtained are shown in table S1.

This approach ensured that all models resulted in similar proportion of first correct decisions, see Figure S2B. The proportion of responses ‘right’ 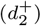 in the second decision are plotted in Figure S2C. As expected, they show a pattern of marked overconfidence: all sampling models tended to respond 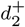 more often than the optimal model, despite similar accuracy in the first decision. The bias is larger for models with smaller number of samples, and decreases approaching the optimal Bayesian model as *n* increases. Importantly, this bias is incompatible with the observed behavioral data, which showed on average a marked under-confidence bias (see Main text, Results).

**Table S1:** Estimated values of *σ* (and their standard errors) that result in similar performance as the Bayesian model with *σ* =1.

| n. of samples | $\sigma$ | se |
| --- | --- | --- |
| 2 | 0.74 | 0.06 |
| 3 | 0.83 | 0.06 |
| 4 | 0.84 | 0.06 |
| 5 | 0.88 | 0.07 |
| 6 | 0.89 | 0.07 |
| 7 | 0.91 | 0.07 |
| 8 | 0.91 | 0.07 |
| 9 | 0.93 | 0.07 |

**Table S2:** Maximum likelihood parameter estimates. For each experiment, condition and model the table shows the mean and the standard deviation (within parentheses) of the parameter estimates across participants.

| Condition | Experiment | Model | $m$ | $w_1$ | $\theta_2$ | $w_2$ | $\theta_3$ | $w_3$ | $\theta_4$ |
| --- | --- | --- | --- | --- | --- | --- | --- | --- | --- |
| D-D | correlated-pairs | biased-Bayesian | 2.06 (1.58) |  |  |  |  |  |  |
| D-O | correlated-pairs | biased-Bayesian | 2.13 (3.01) |  |  |  |  |  |  |
| O-O | correlated-pairs | biased-Bayesian | 3.01 (2.47) |  |  |  |  |  |  |
| D-D | random-pairs | biased-Bayesian | 1.27 (0.68) |  |  |  |  |  |  |
| O-O | random-pairs | biased-Bayesian | 5.97 (3.75) |  |  |  |  |  |  |
| D-D | correlated-pairs | fixed-bias |  |  | -0.67 (0.24) |  |  |  |  |
| D-O | correlated-pairs | fixed-bias |  |  | -0.9 (0.94) |  |  |  |  |
| O-O | correlated-pairs | fixed-bias |  |  | -0.63 (0.3) |  |  |  |  |
| D-D | random-pairs | fixed-bias |  |  | -0.84 (0.46) |  |  |  |  |
| O-O | random-pairs | fixed-bias |  |  | -0.23 (0.22) |  |  |  |  |
| D-D | correlated-pairs | discrete (2 lvl) |  | 0.81 (0.66) | -2.11 (1.52) |  |  |  |  |
| D-O | correlated-pairs | discrete (2 lvl) |  | 0.64 (0.45) | -1.55 (1.12) |  |  |  |  |
| O-O | correlated-pairs | discrete (2 lvl) |  | 1.06 (0.97) | -2.34 (1.77) |  |  |  |  |
| D-D | random-pairs | discrete (2 lvl) |  | 0.55 (0.32) | -1.66 (0.86) |  |  |  |  |
| O-O | random-pairs | discrete (2 lvl) |  | 1.63 (0.47) | -2.85 (2.28) |  |  |  |  |
| D-D | correlated-pairs | discrete (3 lvl) |  | 0.58 (0.56) | -0.73 (0.71) | 0.56 (0.54) | -2.11 (2.1) |  |  |
| D-O | correlated-pairs | discrete (3 lvl) |  | 0.43 (0.5) | -0.83 (0.71) | 1.04 (0.83) | -1.56 (1.39) |  |  |
| O-O | correlated-pairs | discrete (3 lvl) |  | 0.61 (0.53) | -1.04 (1.31) | 0.92 (1.35) | -1.59 (1.68) |  |  |
| D-D | random-pairs | discrete (3 lvl) |  | 0.44 (0.4) | -0.76 (0.27) | 0.47 (0.73) | -1.04 (0.74) |  |  |
| O-O | random-pairs | discrete (3 lvl) |  | 1.08 (0.27) | -0.07 (0.16) | 0.69 (0.43) | -2.78 (2.27) |  |  |
| D-D | correlated-pairs | discrete (4 lvl) |  | 0.53 (0.55) | -0.61 (0.51) | 0.54 (0.5) | -1.06 (0.9) | 0.43 (0.47) | -1.42 (1.53) |
| D-O | correlated-pairs | discrete (4 lvl) |  | 0.42 (0.51) | -0.84 (0.59) | 0.76 (0.8) | -0.76 (1.07) | 0.7 (0.67) | -1.08 (1.12) |
| O-O | correlated-pairs | discrete (4 lvl) |  | 0.94 (0.91) | -0.67 (0.58) | 0.8 (1.62) | -1.46 (1.72) | 0.44 (0.66) | -1.05 (1.63) |
| D-D | random-pairs | discrete (4 lvl) |  | 0.43 (0.41) | -0.67 (0.24) | 0.4 (0.69) | -0.93 (0.78) | 1.3 (1.52) | -0.6 (0.65) |
| O-O | random-pairs | discrete (4 lvl) |  | 1.31 (0.44) | -0.64 (1.44) | 1.11 (0.9) | -1.62 (1.86) | 0.39 (0.55) | -1.65 (2.18) |

**Figure S1:**
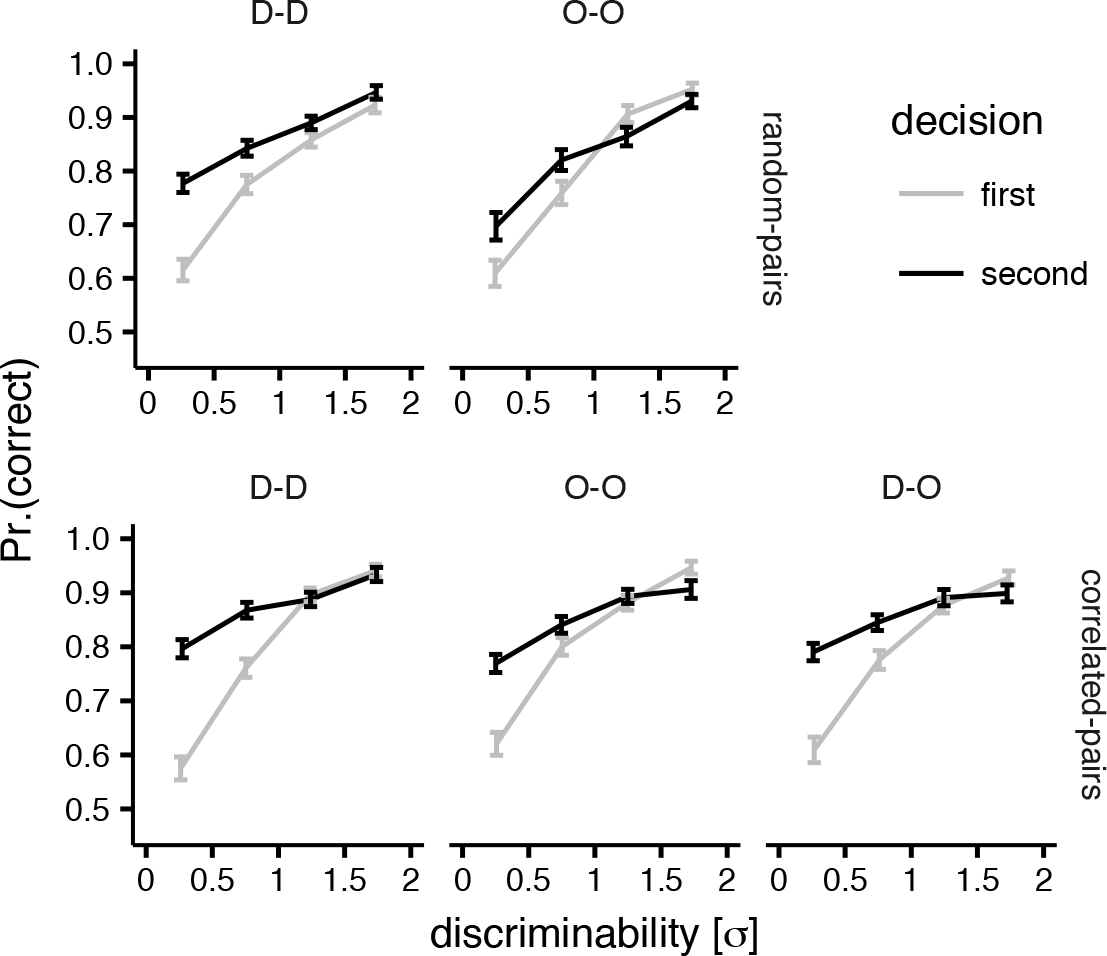
Performance. Same conventions as Figure 1B.

**Figure S2:**
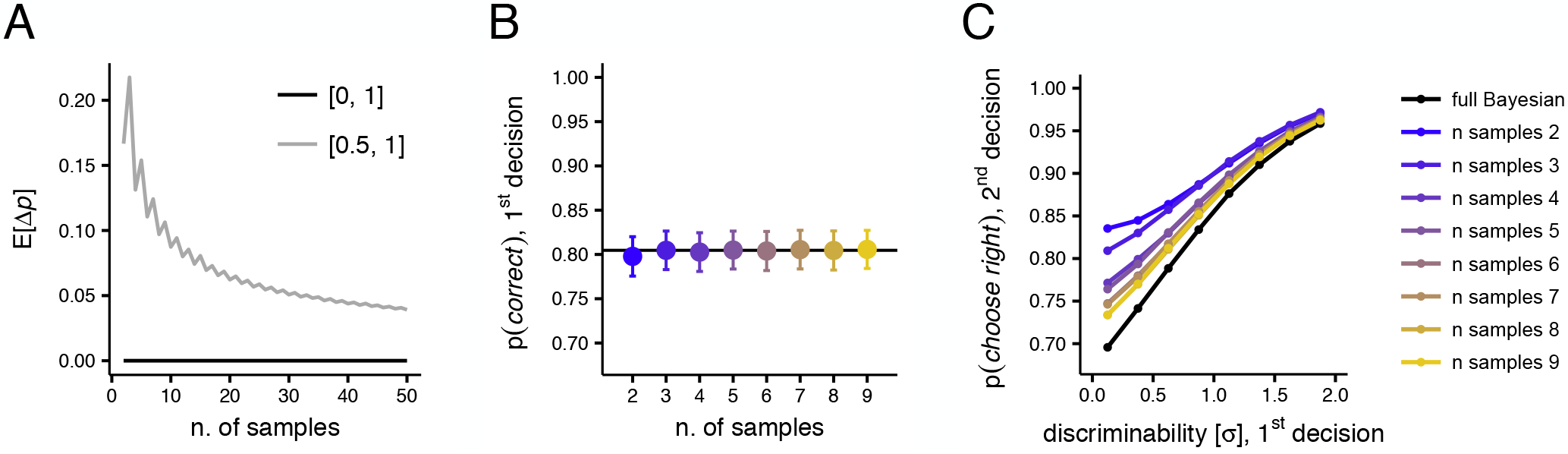
Sampling-based approximation of the Bayesian model. Expected probability estimation bias Δ*p* plotted as a function of the number of samples when the probability *p* varies randomly (uniformly) either in the range [0.5, 1], plotted in grey, or in the range [0, 1] (**A**). It can be seen that when *p* varies within the range of confidence, from chance to certainty [0.5, 1] the predominant bias is one of over-estimation (because Δ*p* is always positive, grey line). Only when *p* varis over the whole domain of probability, [0, 1], the expected bias is on average zero and over-estimation and under-estimation are equally likely (black line). Proportion of correct first decision in the sampling model as a function of the number of samples *n*; the horizontal black line indicates the average performance of the full Bayesian model; error bars represents SEM (**B**). Proportions of responses ‘right’ 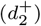 in the second decision as a function of the difficulty of the first decision, as predicted by the full Bayesian model (black line) and the fixed-*n* Bayesian sampler model (**C**).

**Figure S3:**
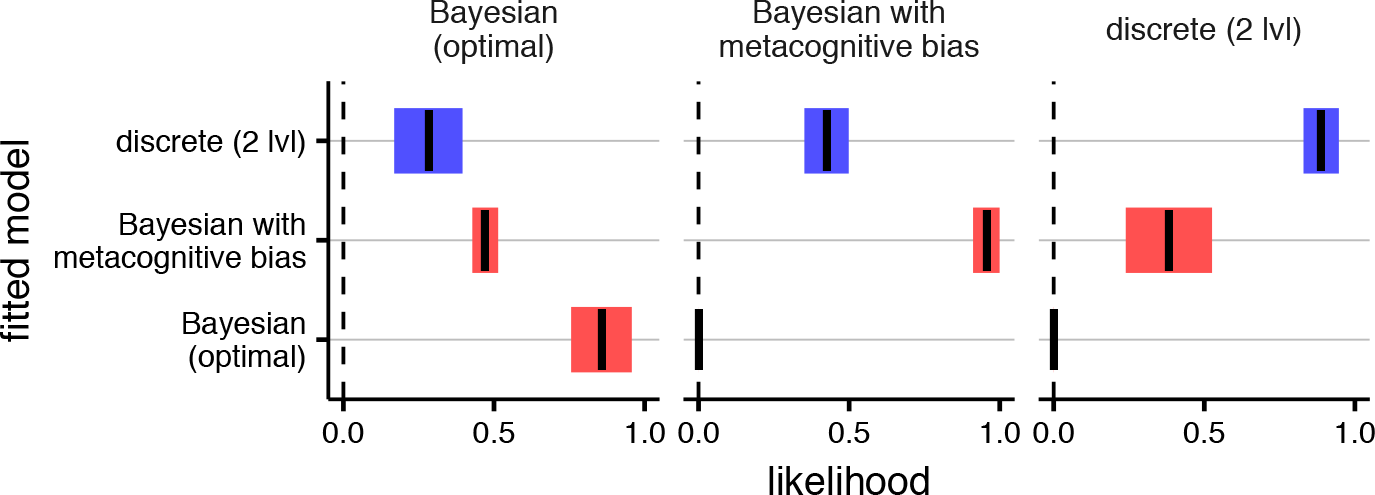
Model recovery analysis. In order to ensure that the models were distinguishable, and to verify that their implementation did not contain errors, we performed a model recovery analysis for our three main computational models: Bayesian (optimal), biased-Bayesian and discrete (2 lvl.). We generated synthetic data (10 simulated observes, for 1000 trials each), with parameters randomly generated from a Gaussian distribution with mean and standard deviations set to the mean and standard deviation of the parameters fitted to our empirical data. Each panel indicate a different generative model, while different lines represent the mean and bootstrapped standard error of the models fit to the synthetic dataset. In each case the model with the highest relative likelihood is the one that generated the data, indicating that the model are correctly recovered and confirm that they are distinguishable.

**Figure S4:**
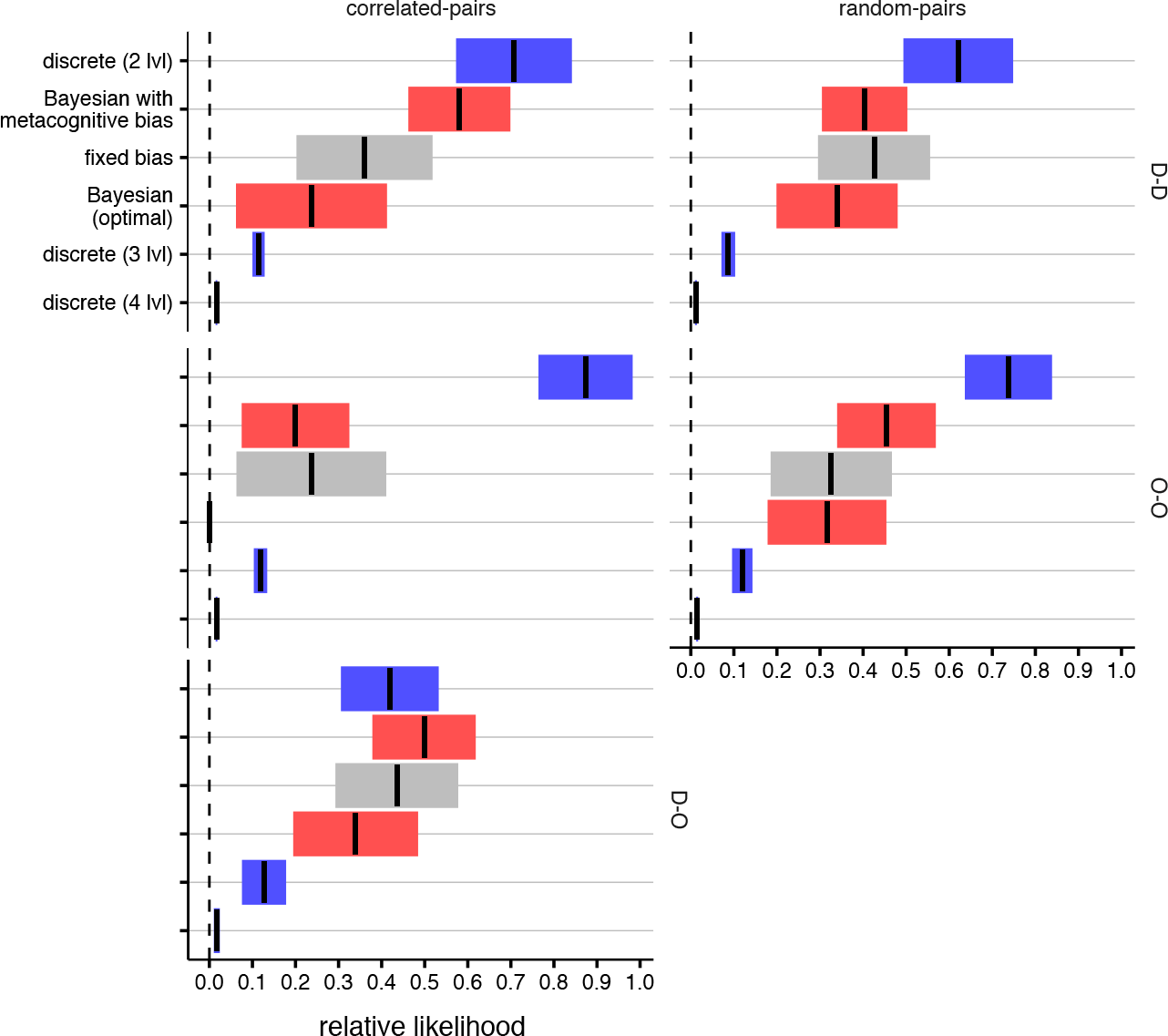
Model likelihoods split according to condition and experiment. Same conventions as Figure 3 in the main text.

## Footnotes

1 Since observers are, overall, more frequently correct than wrong in the first decision (observers made ≈ 80% correct first decisions), the ideal Bayesian model attains an overall better performance (based on the totality of trials).

2 Our analysis indicates that the increase in log-likelihood obtained by adding more than two confidence levels is small and does not compensate the parallel increase in model complexity (number of free parameters), suggesting overfitting.

